# Effects of kinship correction on inflation of genetic interaction statistics in commonly used mouse populations

**DOI:** 10.1101/2021.02.14.431191

**Authors:** Anna L. Tyler, Baha El Kassaby, Georgi Kolishovski, Jake Emerson, Ann Wells, J. Matthew Mahoney, Gregory W. Carter

## Abstract

It is well understood that variation in relatedness among individuals, or kinship, can lead to false genetic associations. Multiple methods have been developed to adjust for kinship while maintaining power to detect true associations. However, relatively unstudied, are the effects of kinship on genetic interaction test statistics. Here we performed a survey of kinship effects on studies of six commonly used mouse populations. We measured inflation of main effect test statistics, genetic interaction test statistics, and interaction test statistics reparametrized by the Combined Analysis of Pleiotropy and Epistasis (CAPE). We also performed linear mixed model (LMM) kinship corrections using two types of kinship matrix: an overall kinship matrix calculated from the full set of genotyped markers, and a reduced kinship matrix, which left out markers on the chromosome(s) being tested. We found that test statistic inflation varied across populations and was driven largely by linkage disequilibrium. In contrast, there was no observable inflation in the genetic interaction test statistics. CAPE statistics were inflated at a level in between that of the main effects and the interaction effects. The overall kinship matrix overcorrected the inflation of main effect statistics relative to the reduced kinship matrix. The two types of kinship matrices had similar effects on the interaction statistics and CAPE statistics, although the overall kinship matrix trended toward a more severe correction. In conclusion, we recommend using a LMM kinship correction for both main effects and genetic interactions and further recommend that the kinship matrix be calculated from a reduced set of markers in which the chromosomes being tested are omitted from the calculation. This is particularly important in populations with substantial population structure, such as recombinant inbred lines in which genomic replicates are used.

## Introduction

In recent years it has become increasingly common to account for relatedness, or kinship, among individuals in genetic association studies, both in human GWAS [1,2] and in model organism studies [3]. Both population structure and cryptic relatedness can lead to artificial inflation of association statistics leading to false positives and loss of power to detect true positive associations [4–6].

A popular method of kinship correction among mouse geneticists is to model relatedness as a random effect using linear mixed models as described in Kang *et al*. (2008) [7]. This method was originally developed to correct kinship effects on genetic main effects in highly structured mouse populations, such as the hybrid mouse diversity panel (HMDP) [7] or multi-generation populations from advanced intercross lines (AIL) [8]. The effects of kinship corrections on main effects in these types of populations are well studied, and have been shown to dramatically reduce false positive rate (FPR) and increases power to detect true main effects [7,8]. Relatively unstudied, however, are the effects of population structure and relatedness on genetic interaction test statistics.

Genetic interactions, or epistasis, are an important aspect of describing complex traits. Statistical models of complex traits are improved when epistasis is taken into account [9], particularly when considering individuals at the tails of the trait distribution [10]. Thus, epistasis may contribute to missing heritability and poor replication of genetic associations across human populations [11]. Kinship may influence these pairwise effects similarly to main effects. Understanding, and appropriately adjusting for kinship when studying epistasis are important in reducing FPR while still maintaining power to detect epistatic effects, which are often weaker than main effects.

Previous work suggests that kinship inflates interaction test statistics, and that adjusting specifically for epistatatic kinship effects effectively reduces FPR in interaction test statistics and improves modeling of complex traits through genetic interaction networks [12]. We sought to expand upon this work by surveying a range of commonly used mouse mapping populations. Although it is common to apply kinship corrections universally, the effects of these corrections is relatively unstudied across population types. Here we investigated the effects of kinship on interaction statistics in these commonly used populations that sampled a range of relatedness as well as population structure.

In addition to calculating main effects and interaction effects using linear models, we investigated the effects of kinship on genetic interaction coefficients from the Combined Analysis of Pleiotropy and Epistasis (CAPE). We previously developed CAPE to combine information across multiple traits to infer directed genetic interactions [13,14]. This type of epistasis analysis is distinct from standard single-trait epistasis analysis, in that the interactions are inferred for multiple traits simultaneously and are directional. The most recent version of CAPE published on CRAN implements a LMM kinship correction, and here we investigated whether kinship caused inflation of these statistics, and how the kinship correction would affect any observed inflation.

For this survey, we selected six mouse populations that are commonly used for identifying both main effects and interaction effects. The populations represented a range of relatedness as well as population structure. In each population we assessed the degree of inflation of main effect test statistics, and interaction test statistics from linear models, as well as CAPE test statistics. We also implemented a kinship correction to investigate the impact of these corrections on test statistic inflation. We used the LMM method originally described in Kang *et al*. (2008) [7]. This method corrects for both cryptic relatedness and population structure simultaneously, and can handle nearly arbitrary and complex genetic relationships between individuals [3]. This is a potentially useful feature in many complicated mouse populations, such as in multi-generational outbred populations, or in experiments involving recombinant inbred lines (RILs) with genomic replicates.

We calculated two different kinship matrices for these corrections. For one we used the full set of genotyped markers to create an overall kinship matrix. For the other we calculated reduced kinship matrices using only markers on chromosomes not being tested for association. The overall correction has been shown to be overly stringent and can reduce power to detect main effects [8]. However by calculating kinship matrices leaving out the chromosome being tested simultaneously controls FPR and retains power to detect main effects on the omitted chromosome [8]. This method is called leave-one-chromosome-out, or LOCO. We implemented an extension of LOCO for epistatic tests in which we calculate kinship matrices leaving out the pair of chromosomes containing the markers being tested. Of course, if both markers are on the same chromosome, this is the same as LOCO. Here we call this extension leave-two-chromosomes-out, or LTCO. We compared the effect of these two kinship matrices across all test statistics and all populations.

## Materials and Methods

### Data

We examined genomic inflation in previously published data sets representing commonly used mouse populations. We selected these populations to represent a range of relatedness and population structure. The populations were as follows: a reciprocal backcross [15], an F2 intercross [16], a panel of BXD recombinant inbred lines (RILs) [17–19], a cohort of Diversity Outbred mice [10,20], and a cohort from an advanced intercross line (AIL) [21]. We created a sixth study population by averaging over genomic replicates in the RIL. Averaging over replicates in RILs is common practice. It reduces population structure, but also reduces *n* and the power to detect effects [22]. We refer to this population as the RIL with no replicates (RIL-NR).

We expected that the AIL, F2, and backcross populations would have negligible population structure, but may potentially harbor cryptic relatedness, where random differences in recombination led to some pairs of individuals being more highly related than other pairs of individuals. Outbred and RIL populations are more likely to have population structure that may confound genetic association tests. The Outbred population used here included multiple generations of animals, and the RIL population included genetic replicates.

Each data set is described in more detail below:

### Mouse Populations

#### Advanced Intercross Lines

This advanced intercross line (AIL) was started by crossing a large (LG/J) mouse with a small (SM/J) mouse [23]. The study population used here are all males derived from the 50th filial generation [21]. Mice were assessed for skeletal and muscular traits at 12 weeks of age [21]. The mice were genotyped at 7187 SNPs. Here we analyzed tibia length (Tibia) and soleus weight (Soleus) in 492 mice.

#### Backcross

This population was generated to investigate gene-environment interactions influencing diabetes and obesity [15]. The diabetes-prone New Zealand Obese (NZO/HlLtJ) mouse was crossed to the diabetes-resistant Non-obese Non-diabetic (NON/ShiLtJ) mouse. The F1 generation was then backcrossed to the NON parent. The study population comprised 204 male mice genotyped at 84 MIT markers. For this study, we selected trygliceride level (TG) and high-density lipoprotein (HDLD) levels. We used cross direction (pgm) as a covariate in all runs.

#### F2

This large F2 intercross was generated to investigate genetic influences on bone density traits in mice [16]. This population carried a fixed *lit* mutation in growth hormone releasing hormone receptor (GHRHR), which arose naturally on the C57Bl6/J (B6) background, and was transferred to the C3H/HeJ (C3H) background. C3H mice with the lit mutation have the same body weight as B6 mice with the lit mutation but have higher bone density. The purpose of this cross was to identify genetic factors that increase bone density in the absences of GHRHR. We used 1095 female mice from this cross. They were genotyped at 100 MIT markers. We analyzed percent body fat (pctFat) and trabecular bone thickness (Tb.Th) here.

#### Outbred

We used a of cohort of Diversity Outbred mice [20], which were derived from eight founder strains: 129S1/SvImJ (129), A/J, CAST/EiJ (CAST), NOD/ShiLtJ (NOD), NZO/HlLtJ (NZO), PWK/PhJ (PWK), and WSB/EiJ (WSB). The CAST, PWK, and WSB strains were recently inbred from wild mouse strains, whereas the other five strains were inbred mostly from pet fancy mice with limited genetic diversity [24]. Across all eight strains there are roughtly 45 million SNPs, and because DO mice are outbred, each carries a unique subset of these SNPs. The systematic mating scheme was designed to limit population structure and relatedness. The DO population we used here included 446 individuals, both male and female. We used only the mice that were fed on a chow diet, eliminating those on a high-fat diet. We used sex as a covariate in all runs. We analyzed the change in blood glucose between 6 and 19 weeks of age (change.urine.glucose) and blood glucose levels at 19 weeks of age (urine.glucose2) in this study.

#### Recombinant Inbred Lines (RIL)

The recombinant inbred lines (RILs) we analyzed here were from the BxD panel of RILs. RILs are generated by crossing two parental strains, breeding the progeny for some number of generations to produce recombinant chromosomes, and then inbreeding to generate stable, inbred genotypes. The result is a panel of inbred mice each with a unique combination of genotypes from the parental strains. BxD were generated from an initial cross between the C57Bl/6J (B) mouse and the DBA/2J (D) mouse. We downloaded data from the Mouse Phenome Database [25] on August 5, 2020.

The data we analyzed were from an experiment investigating the genetics of hippocampal anatomy and spatial learning [17–19]. The data set is called Crusio1. We downloaded all traits related to body weight, radial maze performance, and histopathology.

The BxD panel has been genotyped at 7124 markers across the genome. The genotypes are available from GeneNetwork. [26] We analyzed time to complete the radial maze on the first day of training (task_time_d1) and the number of radial arms entered on day five of training (num_arms_d5) in 452 females.

#### Recombinant Inbred Lines, No Replicates (RIL-NR)

This test used the same RIL data set as described above, but we averaged over individuals of the same strain resulting in 55 individuals. Averaging over replicates in a single strain is common practice. This practice reduces structure in the mapping population, but also reduces power to detect effects [22]. Here we examined how averaging across replicates in a strain affected test statitic inflation. We used only females in this analysis to completely eliminate any duplicated genomes.

### Trait selection

CAPE combines information across multiple traits and requires at least two traits as input. It has been observed previously that body weight and size traits are significantly correlated with the proportion of New Zealand Obese (NZO) genotype in an individual (Petr Simicek personal communication). Two of our populations here, the backcross and the outbred populations, include NZO genomes, and using body weight traits in these populations could lead to increased test statistic inflation due to high levels of polygenicity. To reduce this effect, we selected traits from each population that minimized the correlation with the first principal component of the kinship matrix (Supp. Fig. 4, and Supp. Table 7).

### Kinship Matrix Calculation

We use the R package qtl2 [27] to calculate the kinship matrix as described in Kang *et al*. (2008) [7]. This method calculates a similarity matrix based on measured genotypes. This matrix has been shown to correct confounding population structure effectively and is guaranteed to be positive semidefinite. The kinship matrix is calculated as follows:

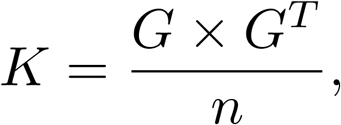

where *G* is the genotype matrix, and n is the number of genotyped markers. For calculating main effects, we use the leave-one-chromosome-out (LOCO) method [8], in which the markers on the chromosome being tested are left out of the kinship matrix calculation. LOCO has been shown to reduce the rate of false negatives relative to use of the overall kinship matrix [8,28]. For each chromosome, we calculated

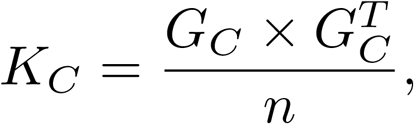

where *G_C_* is the genotype matrix with all markers on chromosome *C* removed. For the pairwise tests, we used the natural extension of LOCO, which we called leave-two-chromosomes-out (LTCO). To calculte the kinship matrix for a pairwise test, we left out the two chromosomes containing the two markers being tested. If both markers were on the same chromosome, we left out only that one chromosome.

#### Fst

To gauge the level of population structure in each population, we calculated the fixation index (*F_ST_*) using the sum of the heterozygosity across all loci π [29]. In the following equation, π_τ_ is the heterozygosity across all populations, and π_S_ is the average heterozygosity across the subpopulations [29].

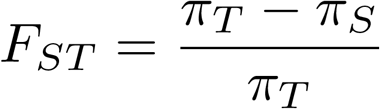

An *F_ST_* of 0 indicates that the population is interbreeding freely, and a value of 1 indicates that subpopulations within the population are genetically isolated. Here, *F_ST_* estimated how structured each mouse population was.

To do this, we converted the kinship matrix for each population to a network using the R package igraph [30]. We used the fast-greedy clustering algorithm [31] in igraph to define subpopulations, which we then used to calculate *F_ST_*.

### Linear Mixed Model Correction

To account for population structure in our association tests, we used a linear mixed model correction as described in [7,32]. Briefly, population corrections account for polygenic effects on the phenotype that are not attributable to the test marker, which cause the assumption of independent prediction errors to fail. To account for correlated errors, Kang *et al*. proposed a mixed-effects model where the residual errors are not independent, but correlated according to a multi-variate Gaussian distribution whose covariance matrix is given by a linear combination of the identity matrix (independent random noise) and a kinship matrix, *K*, which is simply the variance-covariance matrix of the genotypes among individuals. Fitting this model requires identifying the maximum likelihood parameters for the genetic (fixed) effects and the two mixing parameters defining the correlated residual errors. As shown by Lippert *et al*. (2011), this model can be fit rapidly by first factoring K into its spectral decomposition and adjusting the genotypes and phenotypes to align with the residual error structure [32].

The mathematical form of the model allows the fixed effects and the genetic variance to be solved for explicitly as a function of a mixing parameter, which can be optimized using a one-dimensional grid search. We have re-implemented this procedure within CAPE for use with mouse model populations using the code from the R/qtl2 implementation [27].

### Test Statistics from Linear Models

After adjusting for kinship effects, we used single-locus marker regression and pairwise marker regression to derive test statistics in each population. For the single-locus regression, we fit the following model:

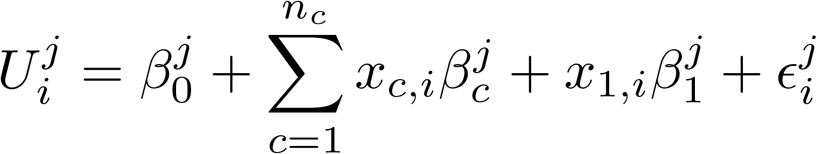

where *U* corresponds to traits, and *∈* is an error term. The index *i* runs from 1 to the number of individuals, and *j* runs from 1 to the number of traits. *x_i_* is the probability of the presence of the alternate allele for individual *i* at locus *j*. We calculated *p* values for each test statistic analytically using a *t* distribution with *n* — 1 degrees of freedom, where n was the number of individuals in the population. We collected main effect test statistics for all traits in each data set.

For the pairwise marker scans, we limited our analysis to two traits. As described below, CAPE requires at least two traits. However, CAPE and pairwise tests in general are computationally intensive, and our ability to run many traits was limited. We fit linear models for each pair of markers and each of the two selected traits as follows:

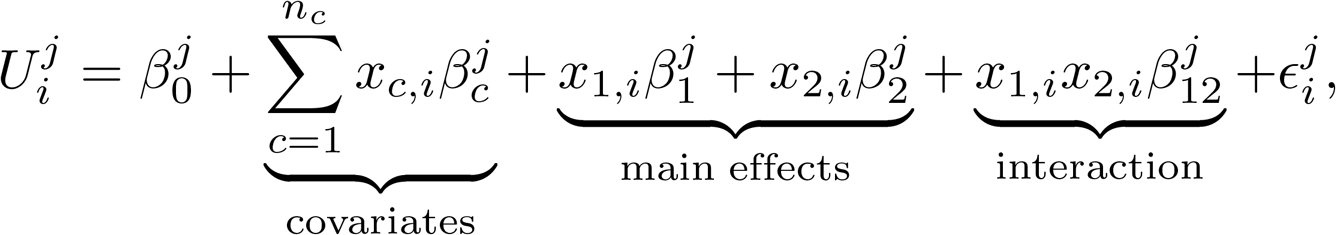

Again, *U* corresponds to traits, and *∈* is an error term. The index *i* runs from 1 to the number of individuals, and *j* runs from 1 to the number of traits. *x_i_* is the probability of the presence of the alternate allele for individual *i* at locus *j*. For the pairwise tests, we calculated empirical *p* values from permutations.

### Combined Analysis of Pleiotropy and Epistasis

Starting with the pairwise linear regression above, we ran the Combined Analysis of Pleiotropy and Epistasis (CAPE) [13,14]. CAPE reparametrizes *β* coefficients from pairwise linear regressions to infer directed influence coefficients between genetic markers. The reparametrization combines information across multiple traits thereby identifying interactions that are consistent across all traits simultaneously. Combining information across traits also allows inference of the direction of the interaction [13,14].

The *β* coefficients from the linear models are redefined in terms of two new *δ* terms, which describe how each marker either enhances or suppresses the activity of the other marker:

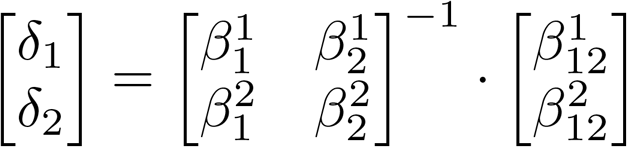

We then translated the *δ* terms into marker-to-marker influence terms:

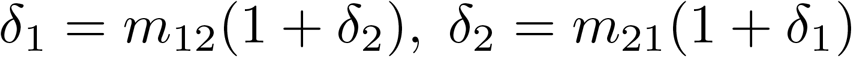

Since matrix inversion can lead to large values with larger standard errors, we performed standard error analysis on the regression parameters, and propagated the errors using a second-order Taylor expansion [14,33]. To calculate *p* values for the directed influence coefficients we performed permutation testing.

### Evaluation of Inflation

To assess test statistic inflation in each population, we ran cape three times, each time collecting the main effect statistics and interaction effect statistics from the linear models, as well as the cape statistics. To estimate the variation in test statistic distributions across sampled populations, we performed Monte-Carlo cross validation [34] by sampling 80% of the individuals over 10 trials.

In each trial, we assessed the inflation of each set of test statistics using *λ* [4]. This inflation factor is the ratio of the median test statistic over the mean of the theoretical distribution. Here we calculated the mean of the chi-square quantiles of 1 — *p* over the theoretical mean of the null, uniform *p* value distribution with one degree of freedom (0.456).

## Results

### Population structure and relatedness varied across populations

We observed varying degrees of relatedness across the populations (Fig. 1). The heatmaps in Fig 1 show how each population was clustered into subpopulations and the relatedness within and among subpopulations. The AIL and F2 populations had negligible structure with no discernible differences in heterozygosity across subpopulations. The Outbred mice were drawn from multiple generations of DO mice, which created subpopulations with slightly higher relatedness than the overall average. The Backcross and RIL had the most substantial structure, with *F_ST_* values more similar to human populations.

**Fig 1.**
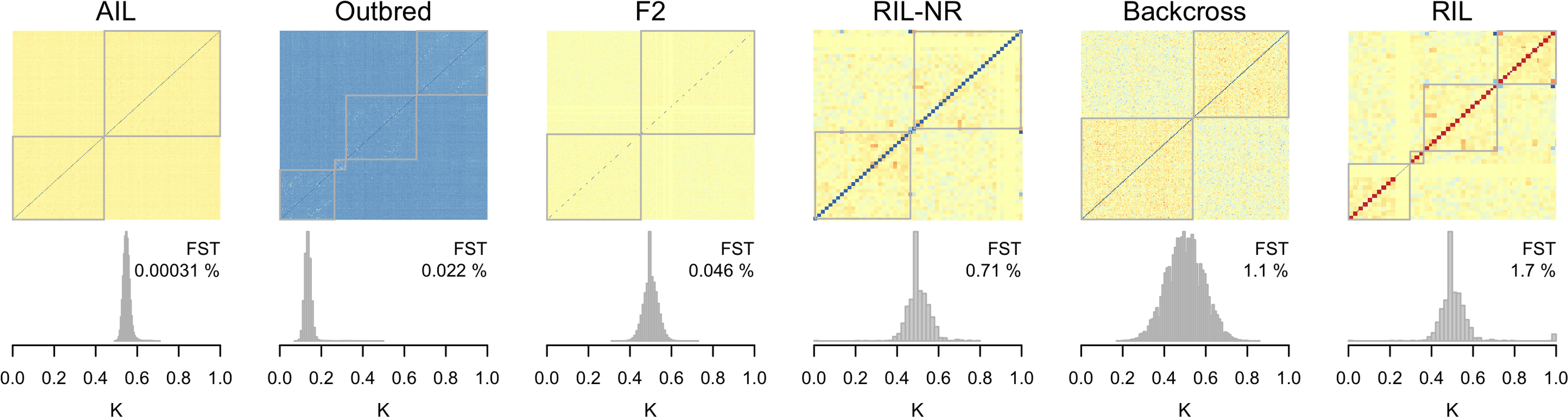
Population structure and relatedness distributions across populations. Each panel shows the population structure of a population as a heatmap in the upper part of the panel. The heatmaps show the overall kinship matrix for each population clustered into subpopulations. Cool colors indicate less relatedness, and warm colors indicate more relatedness. The gray lines indicate the boundaries of subpopulations. The histograms below each heatmap show the distribution of relatedness from the upper triangle of the kinship matrix. Average relatedness varies from the level of cousins in the Outbred animals to slightly higher than the level of siblings in the AIL animals. The *F_ST_* value for each population is shown in the upper right hand corner of each histogram and was calculated as described in the methods. Panels are in order of increasing *F_ST_*.

Independent of the population structure, the populations also had varying degrees of relatedness. On average the Outbred mice were related to each other at a level equivalent to first cousins, which is by design [20], whereas the AIL mice were slightly more related to each other than siblings. The other three populations were all siblings on average, but had differential variation around that mean, with the F2 having a very narrow distribution of relatedness and the Backcross having a wider distribution (Fig. 1).

### Main effect test statistic inflation varied widely across populations

Before running CAPE, we investigated overall trends in test statistic inflation by scanning all traits for main effects using marker regression. This revealed wide variation in test statistic inflation by population (Fig 2). Across all traits, the AIL, Outbred, and RIL-NR populations showed very little inflation. In contrast, the RIL, F2, and Backcross populations showed substantial inflation across most or all traits when no kinship correction was applied (Fig. 2 left-most group). Inflation in the RIL population was corrected by a leave-one-chromosome-out kinship correction (Fig. 2 middle group). The overall kinship correction eliminated inflation in all populations (Fig. 2 right-most group).

**Fig 2.**
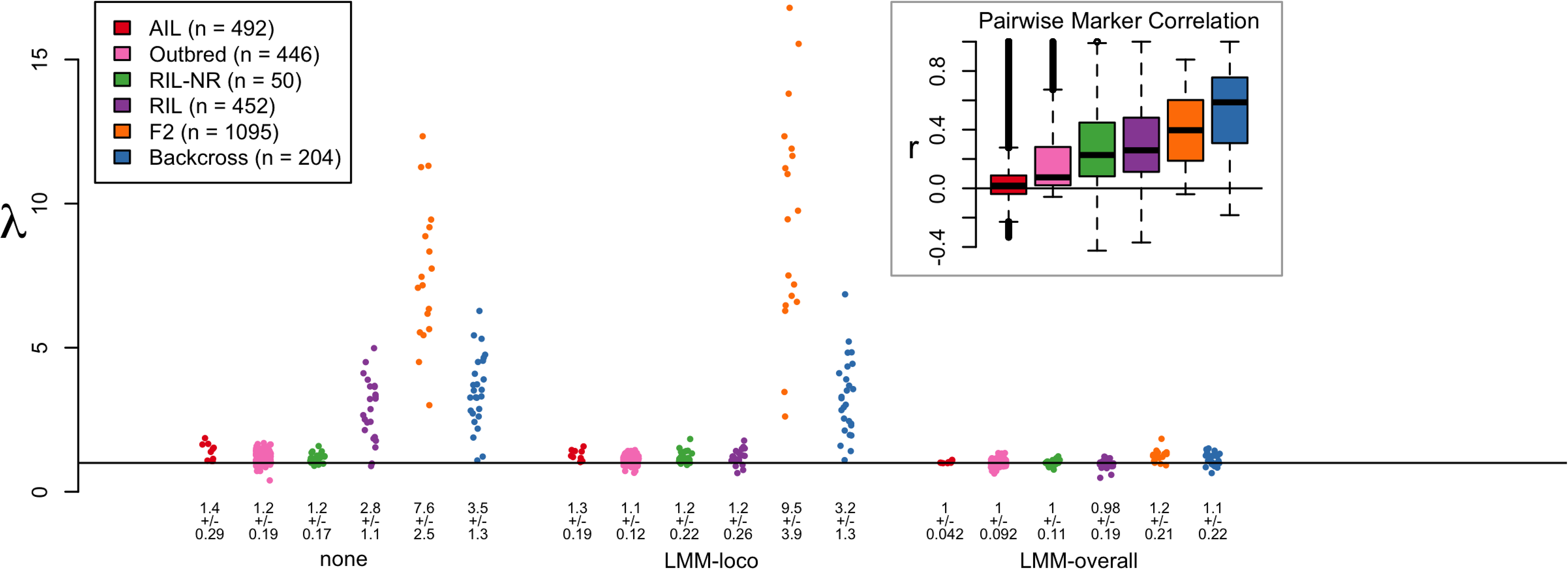
Inflation of test statistics for main effects. Each group of dots shows inflation of main effect statistics across all populations for one of the kinship correction types (none, LMM-loco, or LMM-overall). Each dot represents one trait. The populations are differentiated by color, and are shown in order of increasing LD. The legend shows the correspondance between color and population, as well as the number of individuals in each study. The horizontal line shows λ =1, which indicates no inflation. Numbers below each set of dots indicate the mean and standard deviation of λ for each group. The inset in the top right-hand side of the plot shows the pairwise correlation between markers on the same chromosome for each population, which is a standin for LD. The color of each box identifies which population the data come from. The horizontal line in the boxplot shows *r* = 0. The F2 and Backcross populations, which have the highest LD, also have the highest test statistic inflation. The extreme inflation seen in the F2 population is likely due to a combination of high LD and large n.

### Main effect inflation was correlated with linkage disequilibrium

Linkage disequilibrium (LD) influences test statistic inflation because a single causal SNP within an LD block can inflate the test statistics of all SNPs linked to it. If there are relatively few recombinations in the population, such as in an F2 or backcross, large portions of the genome may be significantly associated with a trait due to linkage alone.

To investigate whether linkage disequilibrium (LD) may be related to the inflation of test statistics in the populations used here, we calculated pairwise Pearson correlations (*r*) between markers on the same chromosome across all chromosomes and all populations. These distributions are shown in the inset in Fig. 2. The two populations with the highest test statistic inflation, the F2 and Backcross populations, also had the highest average LD.

However, although the F2 had lower LD than the backcross, it had substantially greater inflation of test statistics. The F2 also had many more individuals than the backcross, and thus greater power to detect effects. This increase in power combined with high LD could lead to the high levels of inflation seen in the F2. To test this, we subsampled the F2 to the same number of individuals in the backcross and recalculated λ. Reducing n in the F2 also reduced inflation to similar levels seen in the backcross (Supp. Fig 5).

### Kinship corrections reduced inflation differentially across populations

Fig. 3A shows a more detailed view of test statistic inflation in the main effect statistics for each population. Each panel shows QQ plots for the *-log_10_*(*p*) for two traits against the theoretical null *p* values. The more the points rise above the line *y* = *x*, the stronger the inflation factor *λ*. In the absence of a kinship correction, the F2 and RIL showed strong inflation, the AIL and RIL without replicates showed moderate inflation, and the backcross and Outbred populations showed very minor inflation if any at all (Fig. 3A).

Numeric values are shown in the legends of Fig 3. The RIL (λ = 2.7) was the most affected by inflation, while traits in the Outbred population had mild deflation (λ = 0.82).

The overall kinship correction had a strong effect on inflation across all populations (purple dots in Fig. 3A). The leave-one-chromosome-out (LOCO) correction had varied effects (green dots in Fig. 3A). It provided strong control of inflation in the RIL, but had no effect in the Backcross, F2, or AIL populations.

**Fig 3.**
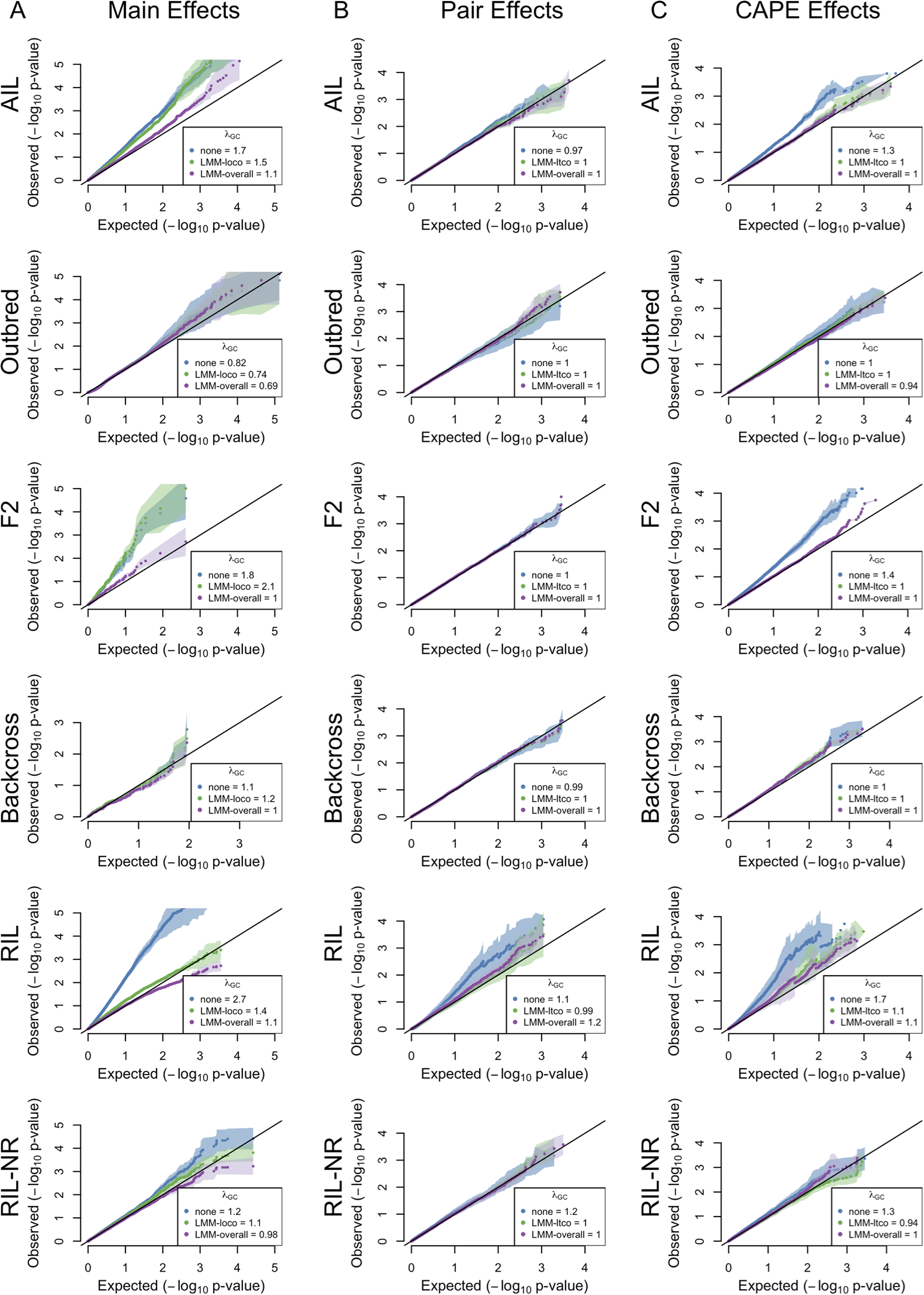
Quantile-quantile (QQ) plots for all test statistics. Each panel shows the QQ plots for one set of statistics across all populations and all correction types. Each row holds the results for a single population. Each column shows one test statistic: (A) QQ plots for main effects. (B) QQ plots for the pairwise test statistics. (C) QQ plots for CAPE statistics. Correction types (none, LMM-loco/ltco, or LMM-overall) are shown in different colors. The *x* axis in each plot shows the theoretical quantiles of the null *p* value distribution, and the *y* axis shows the observed quantiles. Dots show the mean *p* value distribution across 10 rounds of Monte Carlo cross validation, and transparent polygons show the standard deviation. The black line in each plot shows *y* = *x*. The legends show the λ values for each set of statistics.

### Interaction coefficients were largely unaffected by genomic inflation

The *p* values associated with interaction coefficients were almost completely unaffected by inflation (Fig 3B). The only population where inflation appeared to affect the interaction statistics was the RIL population (λ = 1.1). This inflation was reduced by both the leave-two-chromosomes-out (LTCO) and overall kinship corrections.

The LTCO correction appeared to slightly improve power to detect interaction effects in the Outbred population, although this was not evident in the *λ* values (*λ_none_* = 1, vs. *λ_itco_* =1).

### CAPE coefficients were intermediately affected by inflation

The CAPE coefficients influenced by inflation at a level in between that of the main effect statistics and the pairwise statistics (Fig. 3C). The bulk of the inflation was seen in the RIL (λ = 1.7), F2 (λ = 1.4), and backcross populations (λ = 1). The overall and LTCO kinship corrections had remarkably similar effects across all populations.

## Discussion

In this study we examined inflation of main effect and genetic interaction statistics in five mouse mapping populations. We also investigated the effect of kinship corrections on this inflation.

We found large variation in test statistic inflation across populations and across traits. Across populations, the primary driving factors of inflation were LD and population size. Populations with high LD, like the F2 and backcross, had the highest inflation. Between those populations with the highest LD, the number of individuals in the population had a large effect on inflation. High power to detect effects combined with high LD creates hugely inflated test statistics. There was also wide variation in inflation across different traits. We hypothesize that polygenicity may be the primary factor in the variation in inflation across traits within a single population. All else being equal, there will be a preponderance of small *p* values for traits with multiple true positive loci.

Differences in LD cannot explain the difference in inflation between the RIL with replicates, which had substantial inflation, and the RIL without replicates (RIL-NR), which did not. It has been shown previously that including genetic replicates increases power to detect genetic effects [22]. Increase in power alone potentially increases the prevalence of small *p* values; however, genetic relatedness also increases false positive rate (FPR) when strain effects are large relative to individual error [22]. Taken together, these results suggest that including genetic replicates in a RIL study increases power to detect effects, but that a LOCO kinship correction should be done to counterract the increase in FPR caused by the replicates. Here, the LOCO kinship correction substantially reduced inflation in the RIL population without the overcorrection seen with the overall kinship correction (See Figs. 2 and 3A).

The differences of effects between the overall and reduced kinship matrices for the main effects illustrates a couple important points about these two corrections. First, the overall kinship correction reduces power to detect true effects [8]. Indeed, we saw complete elimination of inflation across all populations with this correction. Second, the comparison between the LOCO and overall corrections suggests that the inflation seen in the RIL was primarily due to population structure. The substantial inflation of main effect test statistics in the RIL was reduced by the LOCO correction. However, the LOCO correction did not reduce inflation in the F2 or backcross. These populations had very little structure, and inflation was likely due primarily to LD and polygenicity.

That the overall kinship correction erased all inflation shows how this severe correction can eliminate power to detect true effects. The LOCO correction, however, retains power to detect true effects, while still correcting for relatedness. It should be noted that treating the F2 and Backcross populations as GWAS mapping populations is not really a fair representation, since in practice the markers in these populations would not be treated as independent measurements. However, this exercise illustrates important, albeit dramatic, aspects of test statistic inflation, and how kinship corrections affect test statistics in different situations.

The interaction *β* coefficients did not show any inflation in any population except possibly in the RIL, despite these populations being well powered to detect epistasis. The effects of both kinship corrections were minimal, however, there may have been some minor improvement of power from both corrections in the Outbred population. This complete lack of inflation is in contrast to a previous study in which epistatic test statistics were inflated [12]. There were many differences between this study and the previous study making a direct comparison of results difficult. Ning *et al*. (2018) observed inflation of interaction test statistics in F10 of a mouse advanced intercross line (AIL) [12]. We examined pairwise statistics in later generations of the same AIL. It is unlikely that the reduction in LD or in later generations of the AIL explains the difference in statistic inflation, since the F2 and Backcross in this study had very high LD, and no test statistic inflation. Further, increasing the pairwise marker correlation cutoff to *r* = 0.8 did not change pairwise statistic inflation in any population (data not shown). We performed exhaustive pairwise testing in both the F2 and Backcross, and our F2 was similarly powered to the AIL population in Ning *et al*., suggesting that marker pair sampling and power differences do not sufficiently explain the differences in our observations. However, in the RIL, which was the one population for which there was apparent inflation in pairwise test stastics, both LMM paradigms corrected the inflation. This result is concordant with previous findings that LMM kinship corrections reduce inflation in pairwise test statistics.

In contrast to the interaction coefficients from pairwise linear models, CAPE interaction coefficients did show inflation in some populations. We saw the most inflation in the F2, RIL, and AIL populations. Lambda values were intermediate between those seen for the main effect statistics and the interaction statistics, which we expect given that CAPE interaction coefficients are non-linear combinations of main effect statistics and interaction statistics across multiple traits across populations. We therefore attribute inflation in these CAPE coefficients to propagation of main effect inflation. Indeed, the lambda values of the main effect statistics and CAPE interaction coefficients were positively correlated (Supp. Fig. 6). When there was inflation of CAPE coefficients, both corrections controlled the inflation well. The similarity in effects of the two corrections was somewhat surprising. We predicted that as with LOCO, the LTCO correction would have been less stringent than the overall correction, but this was not what we observed. Extrapolating from the main effect results, test statistic in the RIL should be most subject to inflation derived from kinship. In this population, both kinship matrices controlled inflation well, but the overall correction did trend toward the more severe correction. Although more work needs to be done, these results suggest that using the LTCO kinship matrix for interaction effects may maintain power to detect effects better than the overall matrix.

We conclude that although many experimental mouse populations are created in such a way to minimize population structure, cryptic relatedness and population structure may still increase FPR in these populations for both main effects and genetic interactions. This is particularly true in populations with unusual relatedness patterns, such as RILs with genomic replicates. In all populations, but particularly in those with greater structure, applying a kinship correction reduces FPR. We recommend applying the reduced kinship matrix in which the chromosomes containing the tested markers are left out. These kinship matrices reduce FPR related to population structure with minimal effect on power. The major drawback to implementing these corrections is the computational time they require, particularly for large populations. However, we recommend that any decision to forego a kinship correction should be justified with a full examination of structure in the study population. Simulations were beyond the scope of this project, but could potentially further delineate guidelines for when kinship corrections are necessary, and which types of kinship matrices to use. Such simulations should take LD, polygenicity, and multiple types of population structure into account.

## Acknowledgements

This work was funded by the National Institute of General Medicine grant R01 GM115518.

## Data and Software Availability

All data used in this study and the code used to analyze it are avalable as part of a reproducible workflow on Figshare. The workflow is called Epistasis_and_Kinship_in_Mouse_Populations

CAPE is available at CRAN and on Github at https://github.com/TheJacksonLaboratory/cape.

## Supplemental Figure Legends

**Fig 4.**
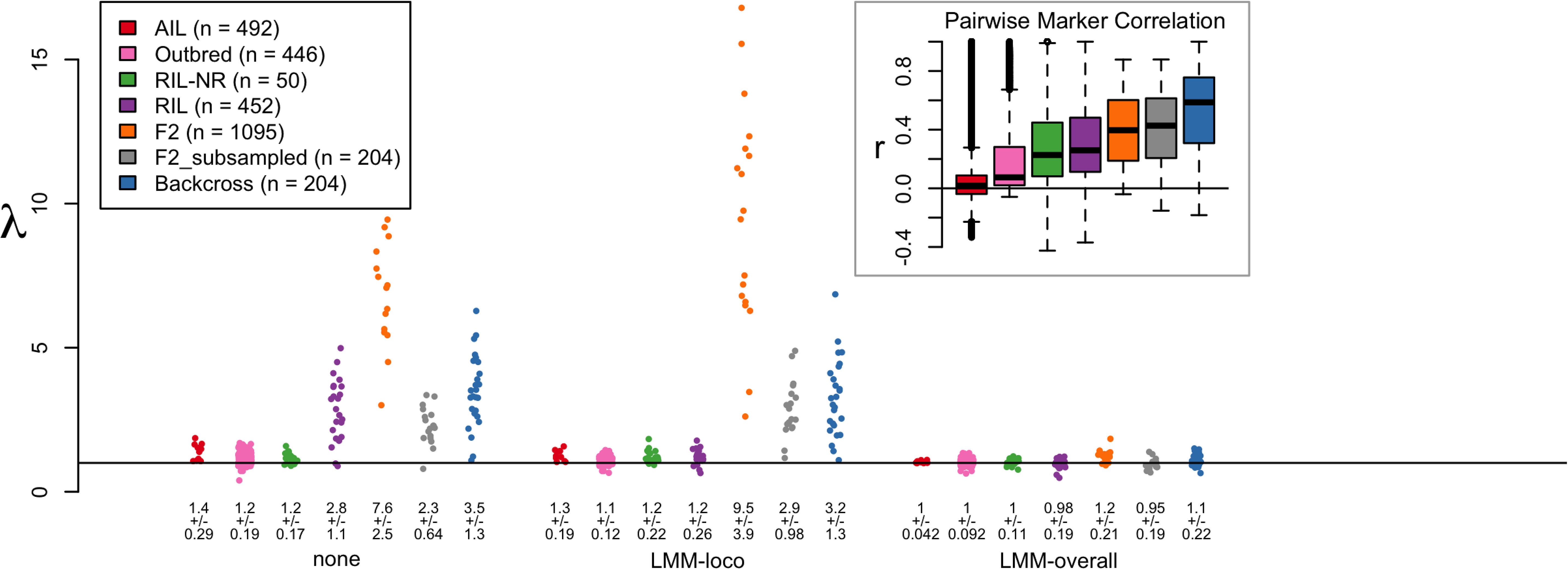
Correlations between traits and the first principal component (PC) of the kinship matrix. Traits with high correlation to the kinship matrix may be highly polygenic and thus be susceptible to test statistic inflation due to many true positives. To reduce this risk, we selected traits with low correlation with the first kinship matrix PC. This figure shows the distribution of correlations between traits and the first kinship PC across populations.

**Fig 5.**
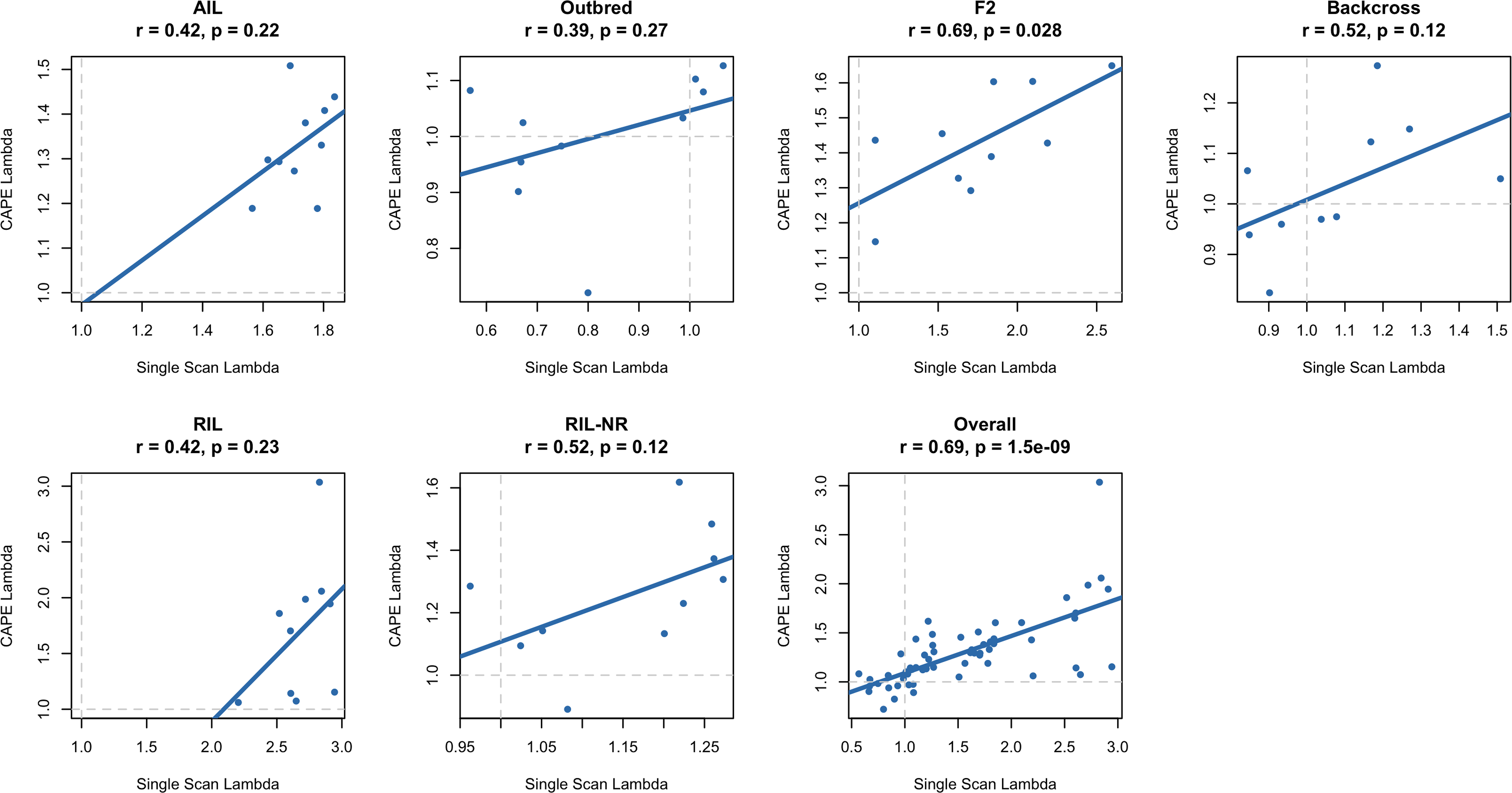
Reducing n reduces inflation. This figure is identical to Fig. 2 except that we have added a column for the F2 that has been subsampled to the same n as the Backcross. This subsampling reduces power to detect effects, and thus reduces inflation to roughly the same level as that seen in the backcross.

**Fig 6.**
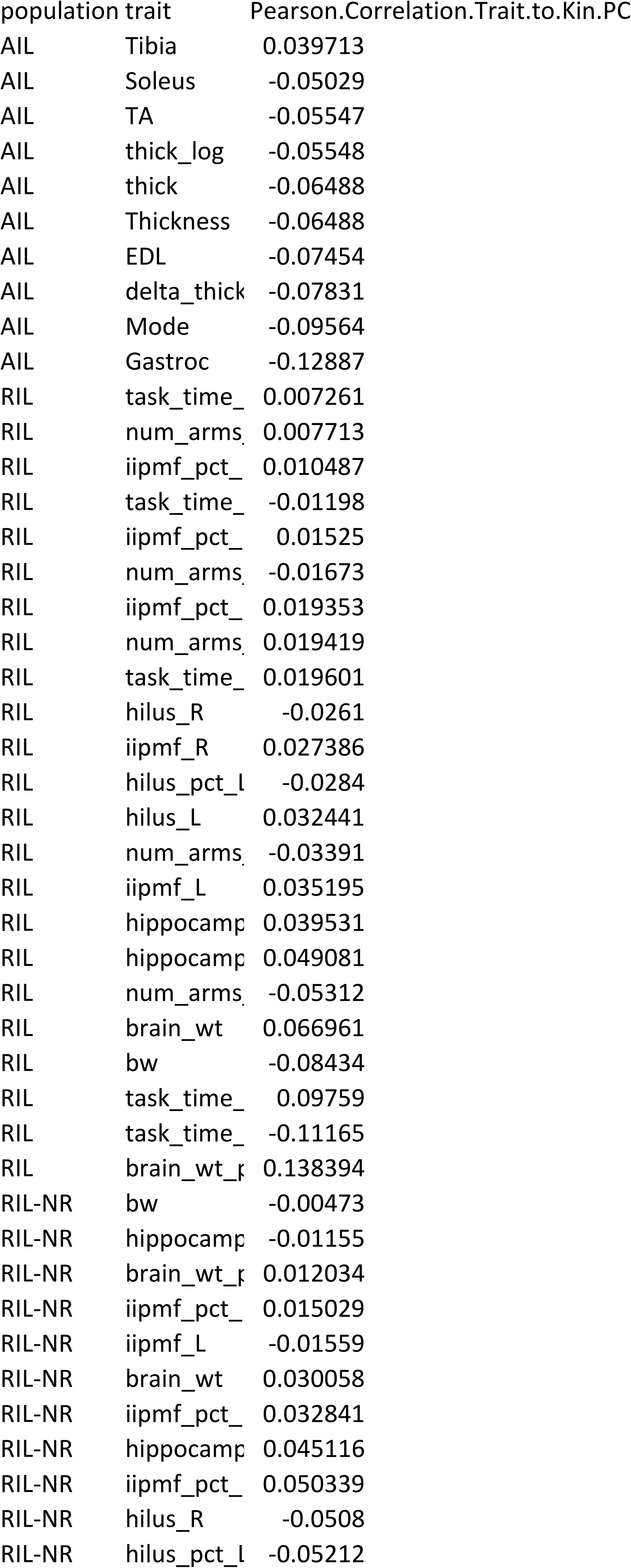
Correlations between lambda values of main effect statistics and CAPE interaction coefficients across populations. Each panel shows the correlation between inflation values for the main effect statistics and CAPE coefficients for a single popopulation. The last panel shows this correlation across all populations. Overall, greater inflation of main effect statistics propagated to greater inflation of CAPE coefficients.

## Supplemental Table Descriptions

**Fig 7.**
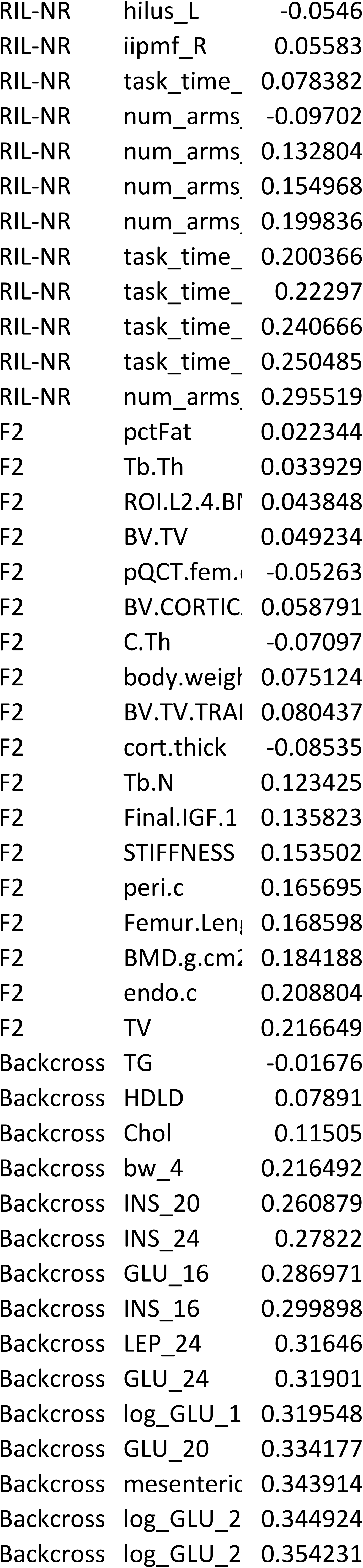
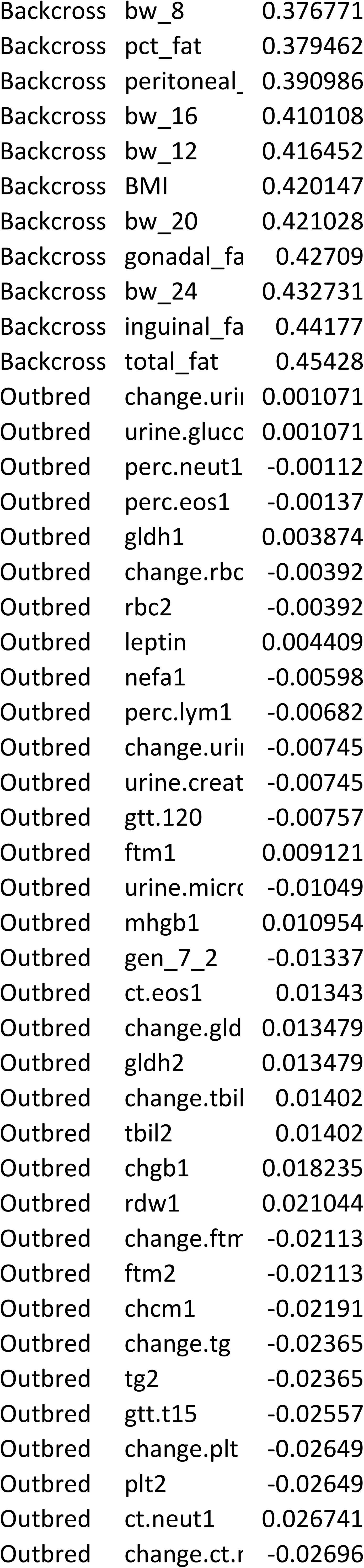
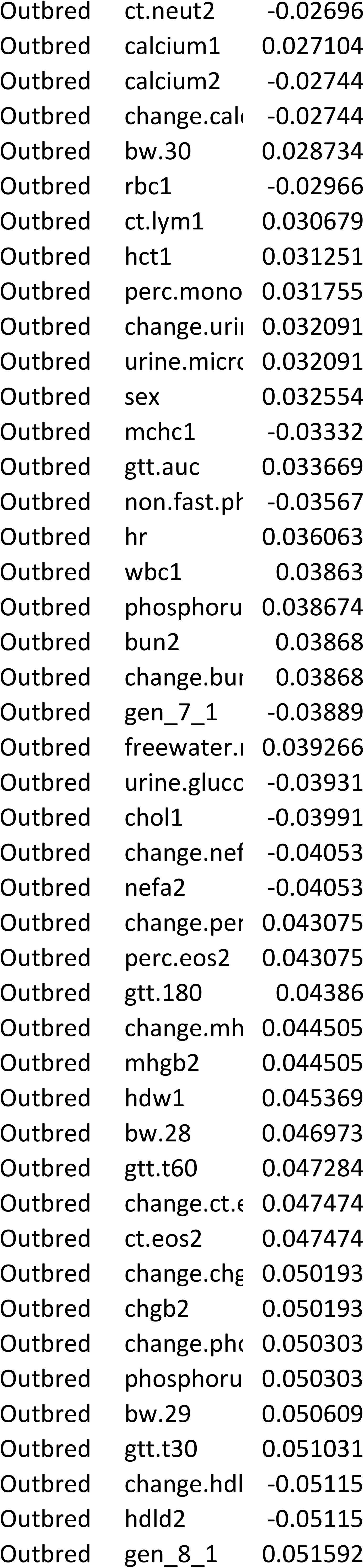
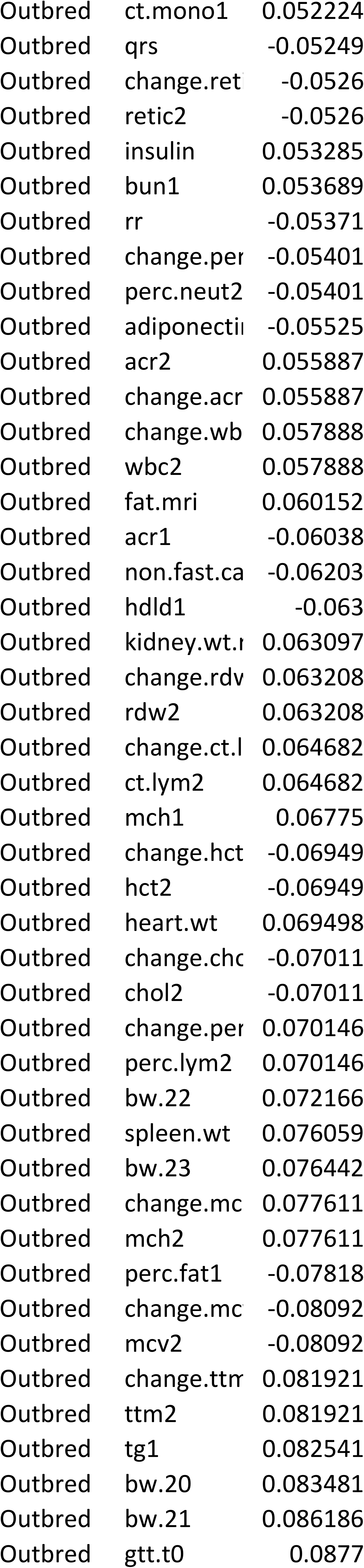
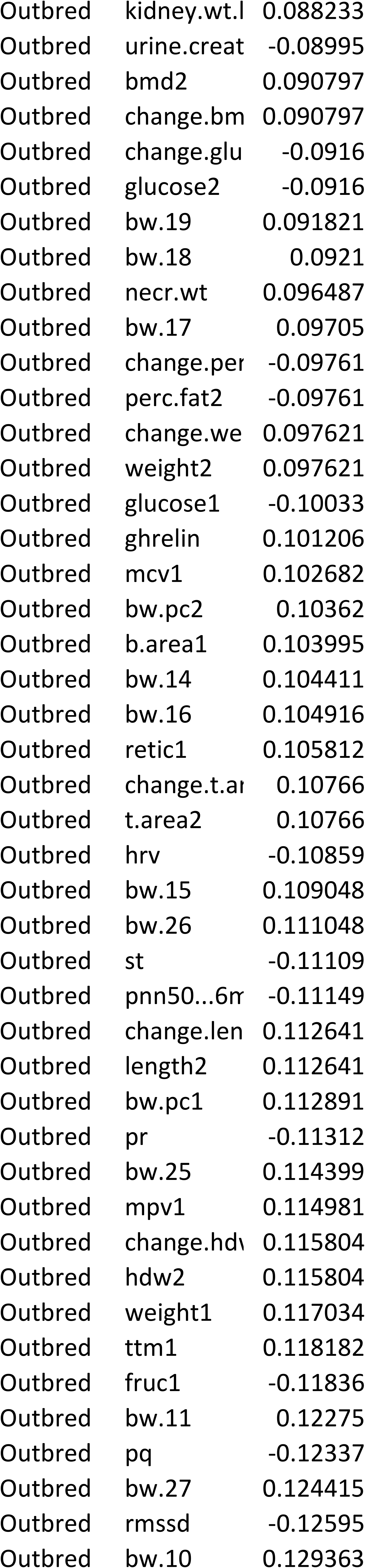
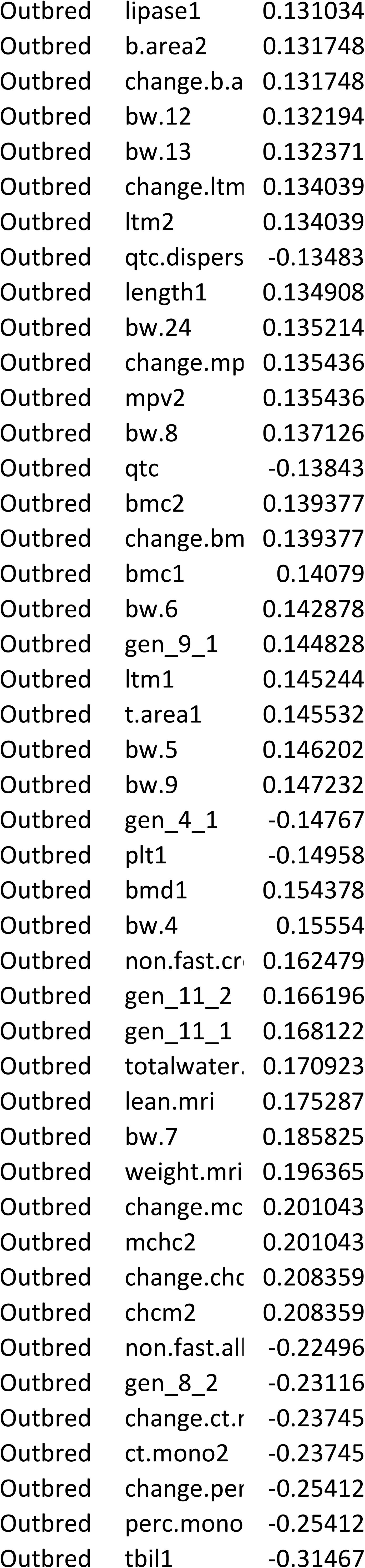
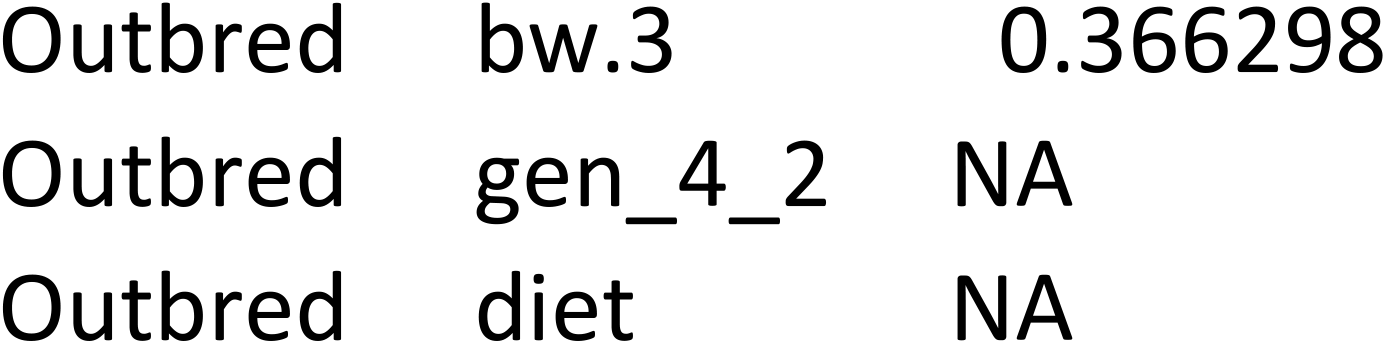
Correlations between traits and the first PC of the kinship matrix.

## References

1. Yao Y, Ochoa A. Testing the effectiveness of principal components in adjusting for relatedness in genetic association studies. BioRxiv. Cold Spring Harbor Laboratory; 2019; 858399.

2. Conomos MP, Miller M, Thornton T. Robust population structure inference and correction in the presence of known or cryptic relatedness. BioRxiv. Cold Spring Harbor Laboratory; 2014; 008276.

3. Sul JH, Martin LS, Eskin E. Population structure in genetic studies: Confounding factors and mixed models. PLoS genetics. Public Library of Science San Francisco, CA USA; 2018;14: e1007309.

4. Devlin B, Roeder K. Genomic control for association studies. Biometrics. Wiley Online Library; 1999;55: 997–1004.

5. Voight BF, Pritchard JK. Confounding from cryptic relatedness in case-control association studies. PLoS Genet. Public Library of Science; 2005;1: e32.

6. Astle W, Balding DJ, others. Population structure and cryptic relatedness in genetic association studies. Statistical Science. Institute of Mathematical Statistics; 2009;24: 451–471.

7. Kang HM, Zaitlen NA, Wade CM, Kirby A, Heckerman D, Daly MJ, et al. Efficient control of population structure in model organism association mapping. Genetics. 2008;178: 1709–1723.

8. Cheng R, Parker CC, Abney M, Palmer AA. Practical considerations regarding the use of genotype and pedigree data to model relatedness in the context of genome-wide association studies. G3: Genes, Genomes, Genetics. G3: Genes, Genomes, Genetics; 2013;3: 1861–1867.

9. Forsberg SKG, Bloom JS, Sadhu MJ, Kruglyak L, Carlborg O. Accounting for genetic interactions improves modeling of individual quantitative trait phenotypes in yeast. Nat Genet. 2017;139: 1455.

10. Tyler AL, Ji B, Gatti DM, Munger SC, Churchill GA, Svenson KL, et al. Epistatic networks jointly influence phenotypes related to metabolic disease and gene expression in Diversity Outbred mice. Geneticsyn. Genetics Soc America; 2017;206: 621–639.

11. Mackay TFC. Epistasis and quantitative traits: using model organisms to study gene-gene interactions. Nature Publishing Group. 2014;15: 22–33.

12. Ning C, Wang D, Kang H, Mrode R, Zhou L, Xu S, et al. A rapid epistatic mixed-model association analysis by linear retransformations of genomic estimated values. Bioinformatics. Oxford University Press; 2018;34: 1817–1825.

13. Tyler AL, Lu W, Hendrick JJ, Philip VM, Carter GW. CAPE: an R package for combined analysis of pleiotropy and epistasis. PLoS Comput Biol. Public Library of Science; 2013;9: e1003270.

14. Carter GW, Hays M, Sherman A, Galitski T. Use of pleiotropy to model genetic interactions in a population. PLoS Genet. Public Library of Science; 2012;8: e1003010.

15. Reifsnyder PC, Churchill G, Leiter EH. Maternal environment and genotype interact to establish diabesity in mice. Genome research. Cold Spring Harbor Lab; 2000;10: 1568–1578.

16. Tyler AL, Donahue LR, Churchill GA, Carter GW. Weak epistasis generally stabilizes phenotypes in a mouse intercross. PLoS genetics. Public Library of Science San Francisco, CA USA; 2016;12: e1005805.

17. Delprato A, Bonheur B, Algeo M-P, Rosay P, Lu L, Williams RW, et al. Systems genetic analysis of hippocampal neuroanatomy and spatial learning in mice. Genes, Brain and Behavior. Wiley Online Library; 2015;14: 591–606.

18. Delprato A, Algeo M-P, Bonheur B, Bubier JA, Lu L, Williams RW, et al. QTL and systems genetics analysis of mouse grooming and behavioral responses to novelty in an open field. Genes, Brain and Behavior. Wiley Online Library; 2017;16: 790–799.

19. Delprato A, Bonheur B, Algeo M-P, Murillo A, Dhawan E, Lu L, et al. A quantitative trait locus on chromosome 1 modulates intermale aggression in mice. Genes, Brain and Behavior. Wiley Online Library; 2018;17: e12469.

20. Svenson KL, Gatti DM, Valdar W, Welsh CE, Cheng R, Chesler EJ, et al. High-resolution genetic mapping using the mouse Diversity Outbred population. Genetics. Genetics Soc America; 2012;190: 437–447.

21. Hernandez Cordero AI, Carbonetto P, Riboni Verri G, Gregory JS, Vandenbergh DJ, P. Gyekis J, et al. Replication and discovery of musculoskeletal qtl s in lg/j and sm/j advanced intercross lines. Physiological reports. Wiley Online Library; 2018;6: e13561.

22. Keele GR, Crouse WL, Kelada SN, Valdar W. Determinants of QTL mapping power in the realized Collaborative Cross. G3: Genes, Genomes, Genetics. G3: Genes, Genomes, Genetics; 2019;9: 1707–1727.

23. Cheverud JM, Routman EJ, Duarte F, Swinderen B van, Cothran K, Perel C. Quantitative trait loci for murine growth. Genetics. Genetics Soc America; 1996;142: 1305–1319.

24. Yang H, Bell TA, Churchill GA, De Villena FP-M. On the subspecific origin of the laboratory mouse. Nature genetics. Nature Publishing Group; 2007;39: 1100–1107.

25. Grubb SC, Bult CJ, Bogue MA. Mouse phenome database. Nucleic acids research. Oxford University Press; 2014;42: D825–D834.

26. Sloan Z, Arends D, Broman KW, Centeno A, Furlotte N, Nijveen H, et al. GeneNetwork: Framework for web-based genetics. Journal of Open Source Software. 2016;1: 25.

27. Broman KW, Gatti DM, Simecek P, Furlotte NA, Prins P, Sen S, et al. R/qtl2: Software for mapping quantitative trait loci with high-dimensional data and multiparent populations. Genetics. Genetics Soc America; 2019;211: 495–502.

28. Gonzales NM, Seo J, Cordero AIH, Pierre CLS, Gregory JS, Distler MG, et al. Genome wide association analysis in a mouse advanced intercross line. Nature communications. Nature Publishing Group; 2018;9: 1–12.

29. Hahn MW. Molecular population genetics. Sinauer Associates New York; 2019.

30. Csardi G, Nepusz T. The igraph software package for complex network research. InterJournal. 2006;Complex Systems: 1695. Available: http://igraph.org

31. Clauset A, Newman ME, Moore C. Finding community structure in very large networks. Physical review E. APS; 2004;70: 066111.

32. Lippert C, Listgarten J, Liu Y, Kadie CM, Davidson RI, Heckerman D. FaST linear mixed models for genome-wide association studies. Nature methods. Nature Publishing Group; 2011;8: 833–835.

33. Bevington PR. Data reduction and error analysis for the physical sciences, ise. New York: McGraw-Hill; 1994.

34. Xu Q-S, Liang Y-Z. Monte carlo cross validation. Chemometrics and Intelligent Laboratory Systems. Elsevier; 2001;56: 1–11.

